# Commensal Microbiota Regulate Renal Gene Expression

**DOI:** 10.1101/2022.12.08.519662

**Authors:** Brittni N. Moore, Jennifer L. Pluznick

## Abstract

The gut microbiome impacts host gene expression not only in the colon, but also at distal sites including liver, white adipose tissue, and spleen. The gut microbiome also influences the kidney and is associated with renal diseases and pathologies; however, a role for the gut microbiome to modulate renal gene expression has not been examined. To determine if microbes modulate renal gene expression, we used whole-organ RNA sequencing (RNA-Seq) to compare gene expression in C57Bl/6 mice that are germ-free (lacking gut microbiota) versus conventionalized (with gut microbiota). 16S sequencing showed that males and females were similarly conventionalized, although Verrucomicrobia was higher in male mice. We find that renal gene expression is differentially regulated in the presence versus absence of microbiota, and that these changes are largely sex-specific. Although microbes also influence gene expression in the liver and large intestine, most differentially expressed genes (DEGs) in the kidney are not similarly regulated in the liver or large intestine. This demonstrates that the influence of the gut microbiota on gene expression is tissue specific. However, a minority of genes (n=4 in males, n=6 in females) were similarly regulated in all three tissues examined, including genes associated with circadian rhythm (*Per1* in males and *Per2* in females) and metal binding (*Mt1 and Mt2* in both males and females). Finally, using a previously published single cell RNA-Seq (scRNA-Seq) dataset, we assigned a subset of DEGs to specific kidney cell types, revealing clustering of DEGs by cell type and/or sex.

**NEW & NOTEWORTHY:** It is unknown whether the microbiome influences host gene expression in the kidney. Here, we utilize an unbiased, bulk RNA-Seq approach to compare gene expression in the kidneys of male and female mice with or without gut microbiota, as well as in liver and large intestine. This report demonstrates that renal gene expression is modulated by the microbiome in a sex- and tissue-specific manner.

## INTRODUCTION

Gut microbiota are influential in maintaining host health(1). Previous reports have highlighted roles for microbial metabolites to alter host signaling pathways(2-5), and recently it has been recognized that microbes also influence the host by modulating gene expression. For example, microbial metabolites alter microRNA expression in *C. elegans*(6), and gut microbes modulate host gene regulation in colonic epithelial cells *in vitro(7)*. Furthermore, germ-free mice (which entirely lack all microbes) have altered gene expression in liver, adipose, and large intestine (8), and germ-free piglets have decreased expression in immune-related genes within oral mucosae(9).

Previous studies have reported a role for gut microbes to affect renal physiology. For example, fermentation of proteins within the gut can lead to the production of excess uremic toxins, such as indoxyl sulfate and *p*-cresyl sulfate(10, 11). These toxins have been implicated in renal disease progression, including chronic kidney disease(10, 11). However, the influence of microbes on renal gene expression has not been explored. To study the relationship between microbiota and renal gene expression, we used bulk RNA sequencing (RNA-Seq) of kidneys from germ-free (GF) and conventionalized (Conv) male and female mice, and examined liver and large intestine transcriptomics for comparison. We find that microbiota influence renal gene expression, and that this influence is largely sex and tissue-specific.

## MATERIALS AND METHODS

### Housing

All animal experiments were approved by the Johns Hopkins Animal Care and Use Committee (accredited by the Association for the Assessment and Accreditation of Laboratory Animal Care). All animals were born in sterile isolators within the Johns Hopkins Germ-Free (GF) Animal Facility. GF mice underwent weekly testing to confirm GF status and were also tested at sacrifice. Testing was performed via gram staining of collected fecal pellets, stool cultures on agar plates, and PCR testing on extracted fecal DNA using primers: 8F (AGA GTT TGA TCC TGG CTC AG) and 1391R (GAC GGG CGG TGW GTR CA).

### Conventionalization

Gut microbiota were introduced to a subset of GF mice (n=10; equal number of males and females) at four weeks of age via an oral gavage using a fecal slurry. The fecal slurry was comprised of mixed stool from age matched, C57Bl/6J animals (males: n=3; females: n=3) who were fed a standard specific pathogen-free (SPF) diet and housed in a standard (not germ-free) Animal Facility. Each GF mouse was inoculated with 120uL of slurry (4% weight by volume into sterile PBS).

### Stool Collection and 16S Microbial Sequencing

Fecal samples from male (n=5) and female (n=5) conventionalized mice were collected at time of sacrifice. Pellets were removed from the distal large intestine, collected on dry ice, and then stored at -80°C. For isolation of bacterial DNA, we used QIAamp Fast DNA Stool Mini Kit (Qiagen, catalog no. 51604), and fecal DNA was isolated following the Qiagen manufacturer protocol. 16S microbial sequencing (2×300bp paired-end reads) was performed using Illumina MiSeq sequencing platform and standard Illumina sequencing primers. The Johns Hopkins University Single Cell and Transcriptomics Core performed library preparation and sequencing, and samples were analyzed by Resphera Biosciences. For analysis, Trimmomatic was used to assess paired-end reads for quality and a minimum length of 200bp(12). FLASH(13) and QIIME(14) were used to merge reads and screen for quality, respectively. Sequences which were spurious hits to the Phix control genome were identified by BLASTIN and removed.

Remaining sequences were stripped of primers, and assessed for chimeras using UCLUST(15) (de novo mode). To screen for contamination in sequencing reads, Bowtie2 was used to filter any mouse-associated contaminants, followed by BLASTN to screen against GreenGenes 16S rRNA database. RDP classifier (80% confidence threshold) was used to identify and filter any mitochondria or chloroplast contamination. High-quality16S rRNA sequences were normalized to 94,000 sequences per sample and assigned to taxonomic lineages for downstream alpha diversity and taxonomic % abundance estimates using Resphera Insight(16). 16S rRNA sequencing reads are available in the National Center for Biotechnology Information database: PRJNA910131.

### Tissue Collection and RNA Isolation

At 8 weeks of age, GF mice (n=10) and conventionalized mice (n=10) were euthanized by CO_2_. All experiments, including sacrifice and tissue collection, were performed during the light cycle between 10:00AM - 1:00PM. Tissues were harvested, placed in TRIzol (Invitrogen, ThermoFisher Scientific, catalog no. 15596026), and stored at -80°C. To collect RNA, tissues were homogenized using a KIMBLE Dounce tissue grinder set (Sigma Aldrich, catalog no.

D8938), and RNA was isolated following the TRIzol protocol. The supernatant from the TRIzol protocol was collected and processed using Qiagen’s RNeasy mini kit. For kidneys, four samples were chosen for each group (male and female germ-free, and male and female conventionalized) based on RNA quality number (RQN) values, and these 16 samples were used for bulk RNA sequencing analysis. For liver and large intestine, 3 samples were chosen from each group based on RQN values.

### Bulk RNA sequencing and analysis

RNASeq (2×150bp paired-end reads) was performed using Illumina NovaSeq sequencing platform. DESeq2 package and output was used to estimate variance-mean dependence of all normalized count data, and to determine differentially expressed genes (DEGs). The total number of genes identified and characterized was 14,949, 13,202, and 15,736 in kidney, liver, and large intestine, respectively. Reads were mapped to mouse reference genome (GRCm39; genecode.vM26). DEGs were defined as having basemean expression > 15 TPM, log_2_ fold change ≥ |0.5|, and p value ≤ 0.5. Data were analyzed by sex as well as microbiota status. RNA-Seq raw counts for all three tissues are available in the National Center for Biotechnology Information database (*kidney:* PRJNA904759; *liver:* PRJNA906570; *large intestine:* PRJNA907282).

To identify intergroup variability of gene expression between samples, we conducted hierarchical clustering using *pheatmap in R* on all three tissues. Raw normalized counts (per kilobases of transcript per 1 million mapped reads; RPKM) for each gene were contained, standard deviation was calculated across each sample for kidney (n=16), liver (n=9), and large intestine (n=9), and a subset of 100 genes were determined to be most variable. An agglomerative approach was used for clustering analysis.

### Single cell RNA-sequencing comparisons

To determine if DEGs clustered to specific kidney cell types, we compared our bulk RNA-Seq data to a previously published single cell RNA-Seq dataset(17). For this dataset, the authors dissected whole kidneys of C57Bl/6J (8-9 week old) male and female mice into 3 zones (cortex, outer medulla, and inner medulla), and sequenced 31,625 cells. The authors assigned the cells to 12 distinct clusters, and the top 50 most differentially expressed genes in each cluster, compared to all other cell types, were identified. To assign genes identified in our bulk RNA-Seq to these distinct clusters, we compared male and female DEGs (Conv versus GF) to the list of genes identified as marker genes by the aforementioned scRNA-Seq study. Log2 fold change values from RNA-Seq data were then assigned to each gene within specific renal cell types and analyzed.

## RESULTS

### Conventionalization is similar in male and female mice

Males and females were conventionalized using a fecal slurry that was an even mix from both male and female donors. To determine if there were any differences in conventionalization in male vs female recipients, we performed 16S rRNA sequencing on stool collected from male Conv and female Conv mice. In parallel, we analyzed the fecal slurry (FS) used for inoculation. Figure 1a shows the relative abundance of top 10 major phyla. Conventionalization was similar in males and females across all phyla with the exception of Verrucomicrobia (1.68% of the gut microbiome in females and 11.17% in males; Fig 1b).

**Figure 1.**
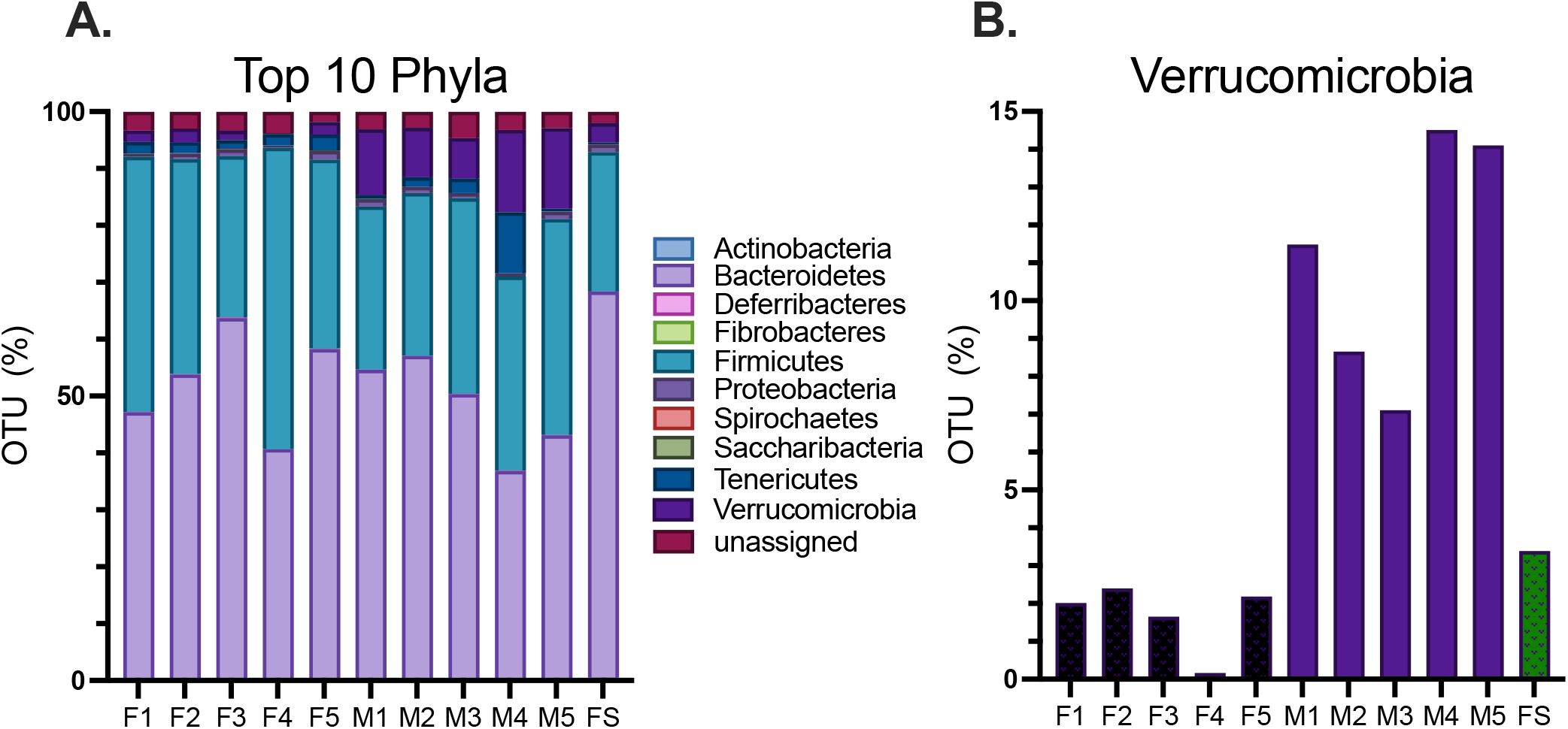
16S sequencing of conventional samples. Fecal samples from conventionalized mice were collected at sacrifice and 16S RNA was sequenced. The fecal slurry (FS) used for inoculation was analyzed in parallel. **A**. Relative abundance of top 10 major phyla shows similar conventionalization in males and females across most phyla. **B**. *Verrucomicrobia* was significantly more abundant (p value= 0.005) among males versus females.

### Both sex and microbial status influence renal gene expression

Principal component (PC) analysis was used to analyze kidney RNA-Seq data (Fig 2). In the kidney, samples cluster by sex (male versus female) on the PC1 axis and by microbiome status (GF versus Conv) on the PC2 axis (Fig 2a). Clustering by sex and by microbiome status was also seen in liver (Fig 2b), but clear clustering by sex and/or microbiota status was not apparent in large intestine samples (Fig 2c).

**Figure 2.**
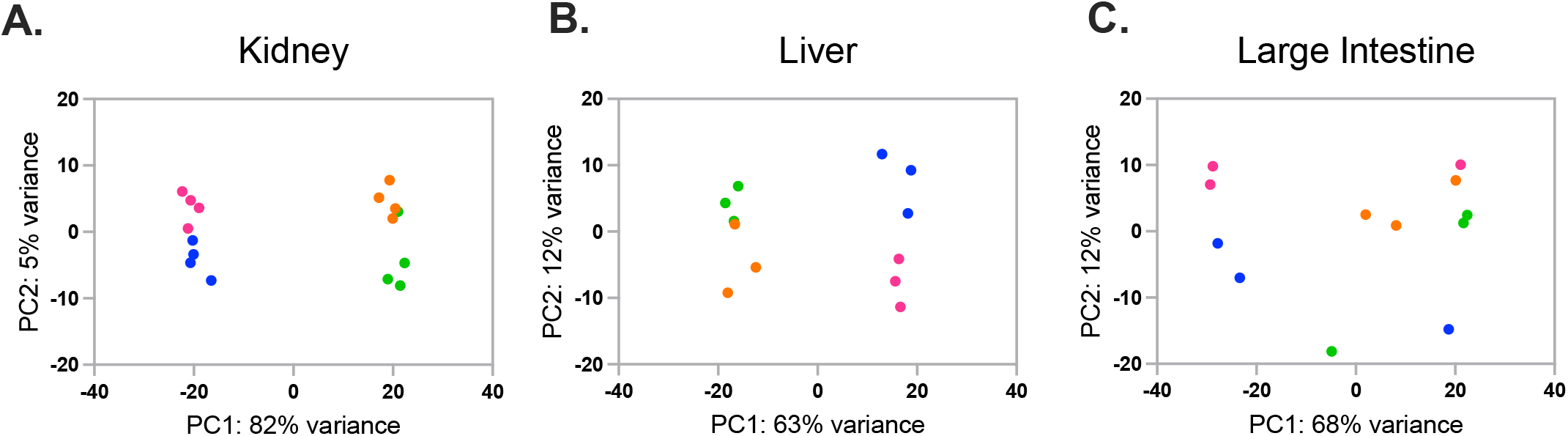
Principal component analysis of RNA-Seq data from kidney, liver and large intestine. Principal component analysis shows the relationship between individual samples from kidney (A), liver (B), and large intestine (C). Data from both kidney and liver cluster by sex along the PC1 axis and microbiota status along the PC2 axis.

In agreement with the PC analysis, unsupervised hierarchical clustering shows that renal gene expression clusters entirely by sex and largely by GF or Conv status (Fig 3a). By comparison, the liver demonstrates similar expression patterns to the kidney (Fig 3b), while in the large intestine, these patterns are less clear (Fig 3c).

**Figure 3.**
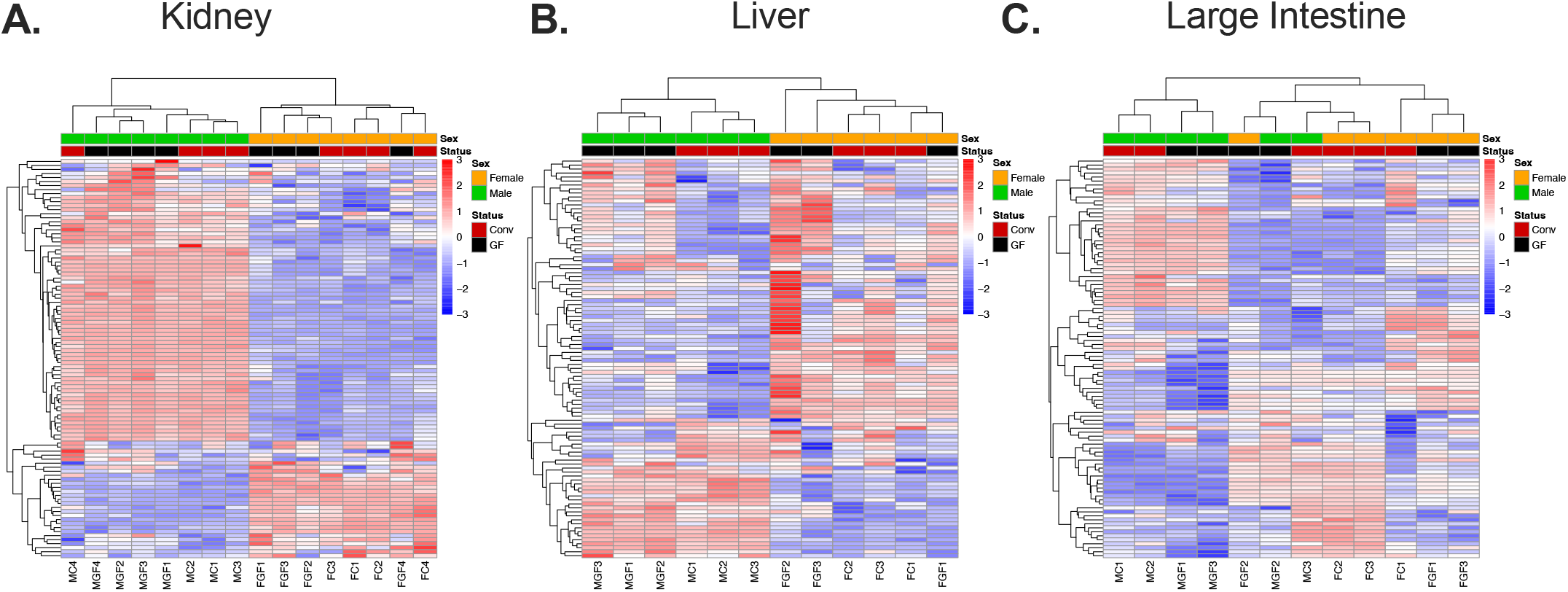
Unsupervised hierarchical clustering of top 100 variable genes in kidney, liver, and large intestine. Raw normalized counts were used to calculate variability across all samples sequenced. **A**. Clustering of top 100 DEGs in kidney shows clear separation by sex. **B**. Sex is dominant factor in clustering of liver DEGs. **C**. Large intestine demonstrates most variability between samples without distinct clustering by sex and/or microbiota status.

### Microbiota-dependent changes in gene expression are unique to the kidney

Genes differentially expressed by microbiome status are displayed in Figure 4. We find a number of genes differentially regulated by microbial status in kidney in both males (340 DEGs) and females (578 DEGs). 51% of these DEGs are upregulated in the presence of gut microbes in females (Fig 4d) and 74% of these DEGs are upregulated in the presence of gut microbes in males (Fig 4i). We also observe a greater number of genes differentially regulated in liver and large intestine compared to kidney, in both females (Fig 4b-d) and males (Fig 4g-i).

**Figure 4.**
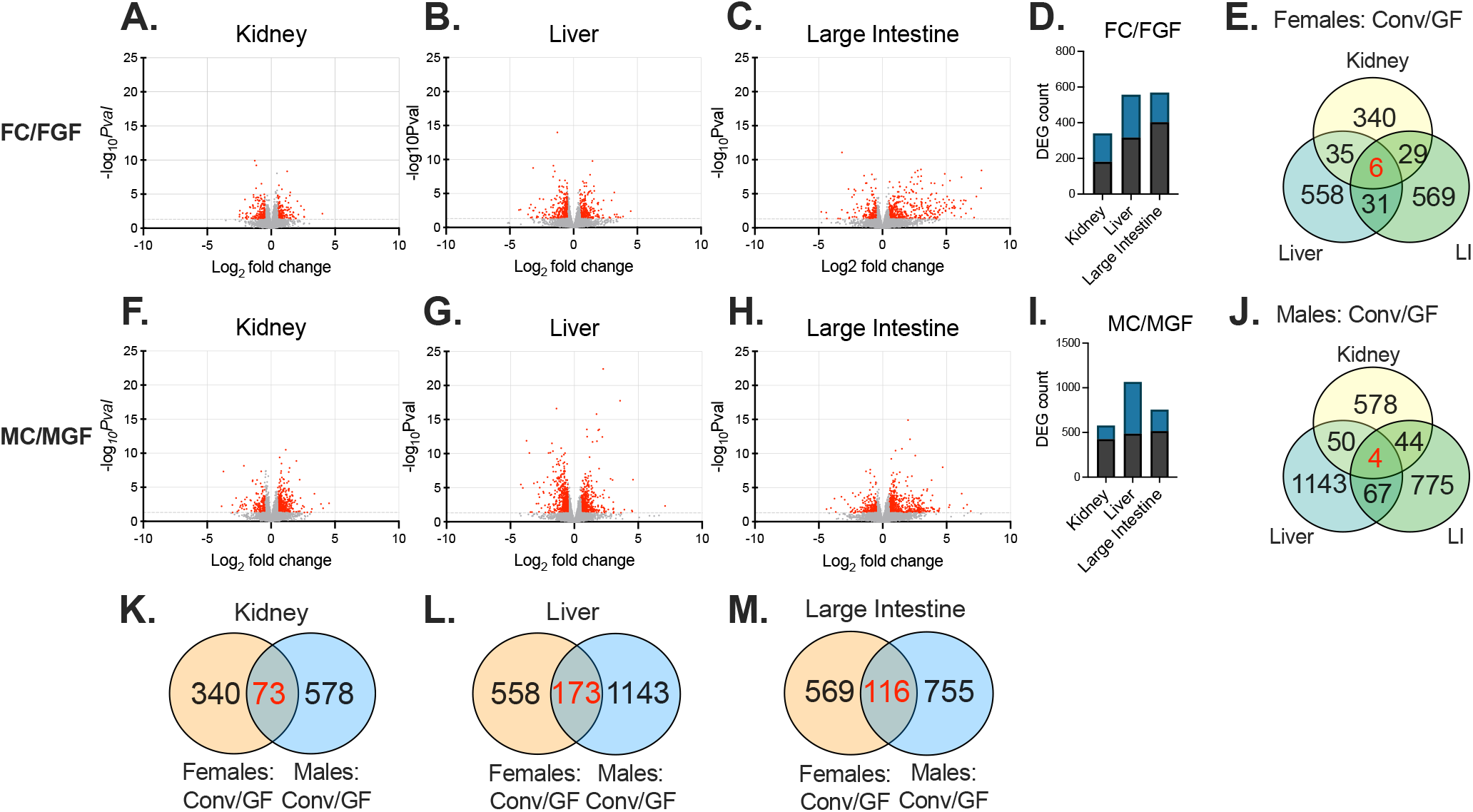
Volcano plot analysis comparing conventional vs germ-free data. Data are shown here separated by both tissue and by sex. The total number of genes identified was 14,949 (kidney), 13,202 (liver), and 15,736 (large intestine). **A-C, F-H**: DEGs are shown in red; genes without significant changes are shown in grey. DEGs are defined as: base mean ≥ 15, |log2 FC| ≥ 0.5, p ≤ 0.05. **D, I:** The total number of DEGs identified in each tissue with respect to male and females (Conv vs GF). Black bars represent the percentage of genes that are upregulated, and blue bars represent the percentage of genes that are downregulated. **E, J:** Venn diagrams show how many DEGs are in common across organs within each sex. **(E)** Six genes co-regulated in females: *Per2, Jchain, Lpl, Extl1, Mt1, and Mt2*. **(J)** Four genes shared in males: *Per1, Mt1, Mt2, and Foxo3*. **K-M:** Venn diagrams compare DEGs altered by gut microbiota in males and females in kidney **(K)**, liver **(L)**, and large intestine **(M)**. Over 90% of genes were either differentially regulated in females or males, and less than 10% of genes commonly regulated in both sexes.

We next investigated if the genes regulated by microbes in the kidney are kidney-specific, or if these genes are globally regulated by microbes. We find that the majority of the genes that changed with microbial status were specific to the kidney (vs liver or large intestine; 80% of DEGs were specific to kidney in females, and 85% of DEGs in males). In fact, only 6 genes changed in all three tissues in females (*Per2, Jchain, Lpl, Extl1, Mt1, and Mt2*), and only 4 genes changed in all three tissues in males (*Per1, Foxo3, Mt1*, and *Mt2*) (Fig 4e,j). *Mt1, Mt2*, and *Per2* are upregulated in GF females compared to Conv females. In contrast, *Lpl, Extl1, and Jchain* are decreased in GF females vs Conv. The four genes which changed across tissues in males were all significantly upregulated in GF compared to Conv.

### Microbes alter renal gene expression in a sexually dimorphic manner

We wondered if microbiota regulate the same genes in males and females. In the kidney, we find that of the 578 DEGs identified in males (Conv versus GF) and the 340 DEGs identified in females, 73 genes are differentially expressed in both sexes (Fig 4*k*). The liver and large intestine exhibited similar sexually dimorphic trends with the majority of DEGs (90% in liver, 89% in large intestine), changing in either males or females, but not both (Fig 4l,m). These analyses demonstrate that the influence of gut microbiota on gene expression is sex specific.

To further explore the role of sex, male-female comparisons are shown in Fig 5. Interestingly, the bulk of genes that are altered based on sex (males versus females, irrespective of microbiota status) occur in the kidney (Fig 5d,i). A subset of genes which are differentially regulated by sex are dependent on the gut microbiota (of the 2536 DEGs in male vs female Conv mice, only 1747 are shared by male vs female GF mice; Fig 5k). Intriguingly, there are over 1000 genes that are differentially regulated by sex in GF kidneys which are not regulated in Conv kidneys (Fig 5k); this may represent sex-specific compensatory changes that occur in the absence of gut microbial signals.

**Figure 5.**
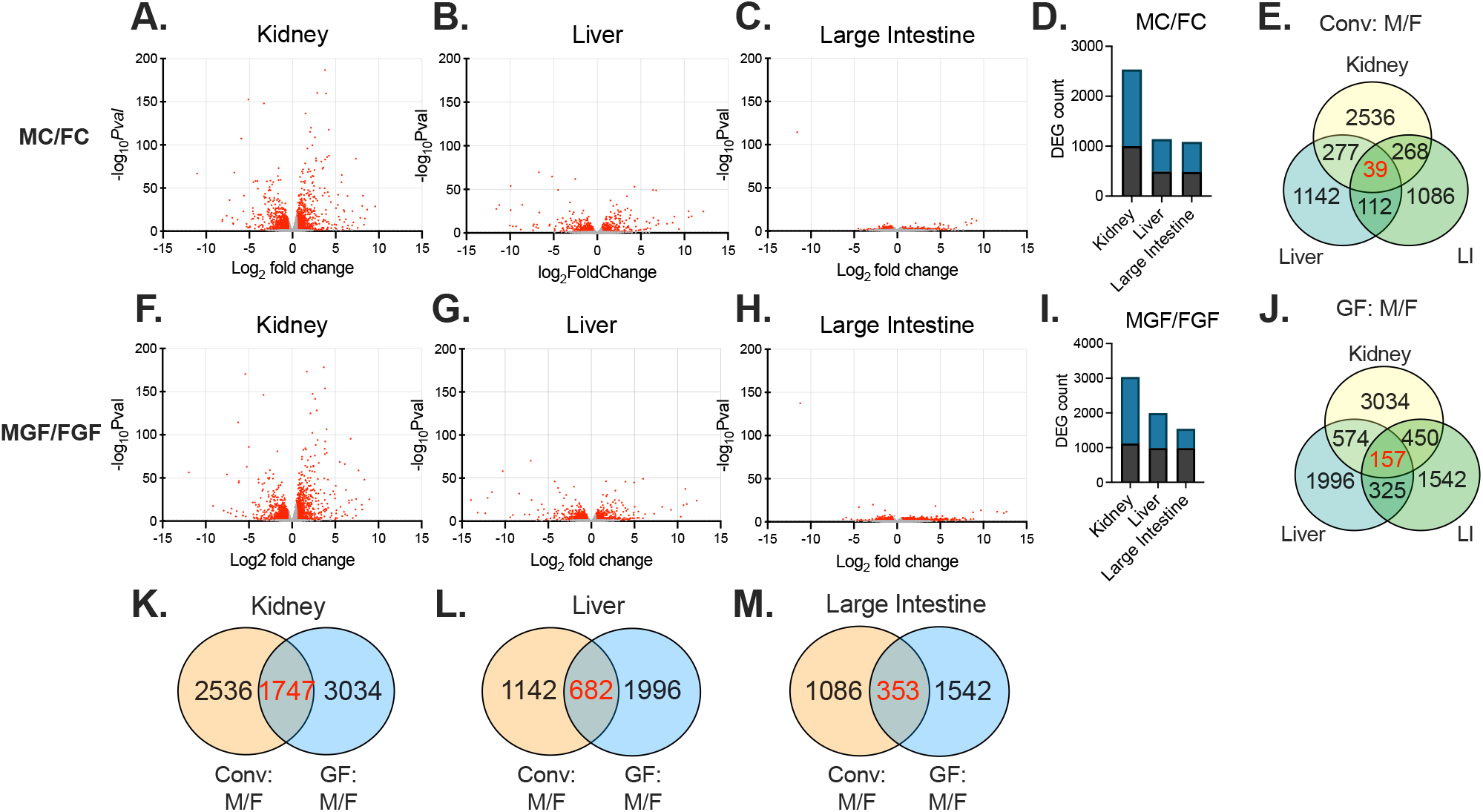
Volcano plot analysis comparing male vs female data. Data are shown here separated by both tissue and by sex. The total number of genes identified was 14,949 (kidney), 13,202 (liver), and 15,736 (large intestine). **A-C, F-H**: DEGs are shown in red; genes without significant changes are shown in grey. DEGs are defined as: base mean ≥ 15, |log2 FC| ≥ 0.5, p ≤ 0.05. **D, I:** The total number of DEGs identified in each tissue with respect to Conv and GF mice (male vs female; M/F). Black bars represent the percentage of genes that are upregulated, and blue bars represent the percentage of genes that are downregulated. **E, J:** Venn diagrams illustrate DEGs differentially regulated by sex which are in common between different organs in Conv and GF mice. **(E)** 39 genes are differentially regulated in kidney, liver, and large intestine in Conv mice. **(J)** 157 genes are shared in GF animals across all three tissues. **K-M:** Venn diagrams illustrate DEGs in common between Conv (male vs female) and GF (male vs female) for each organ. **(K)** 31% of DEGs in the kidney are altered in both Conv and GF animals, **(L)** 22% of genes in liver are differentially expressed based on microbes, and **(M)** less than 15% of genes in large intestine are differentially regulated in Conv vs GF mice, regardless of sex, emphasizing the effect of microbes on gene expression is highly sexually dimorphic.

### Differential gene expression in renal cell types

Next, we compared our data to data from Ransick, et al (17), which performed unbiased single-cell RNA sequencing on kidneys from 2 female and 2 male mice. In this study, “marker genes” of each renal cell type were identified based on the top 50 most differentially expressed genes within each cluster. Comparing our kidney DEGs (Conv versus GF) to genes identified as marker genes in Ransick, et al, we were able to match 176 DEGs from our male data to at least one cell type, and 33 DEGs from our female data to at least one cell type (Table S1,S2). We find that in females, the clusters with the largest magnitude of log 2 fold changes belonged to principal cells (log2fc between +1.73 and -0.54), vascular cells (log2fc between +0.66 and - 1.03), and the proximal tubule (log2fc between +0.57 and -0.96) (Fig 6a). In males, these clusters are NK-myeloid cells (log2fc between +1.69 and -1.99), proximal tubule (log2fc between +1.45 and -0.96), and vascular cells (log2fc between +1.15 and -1.14) (Fig 6b). This implies that there may be commonality in the cell types being regulated by microbes between male and females, even when the genes themselves differ between sexes. Altogether, these data suggest microbes regulate specific genes within various cell types in the kidney.

**Figure 6.**
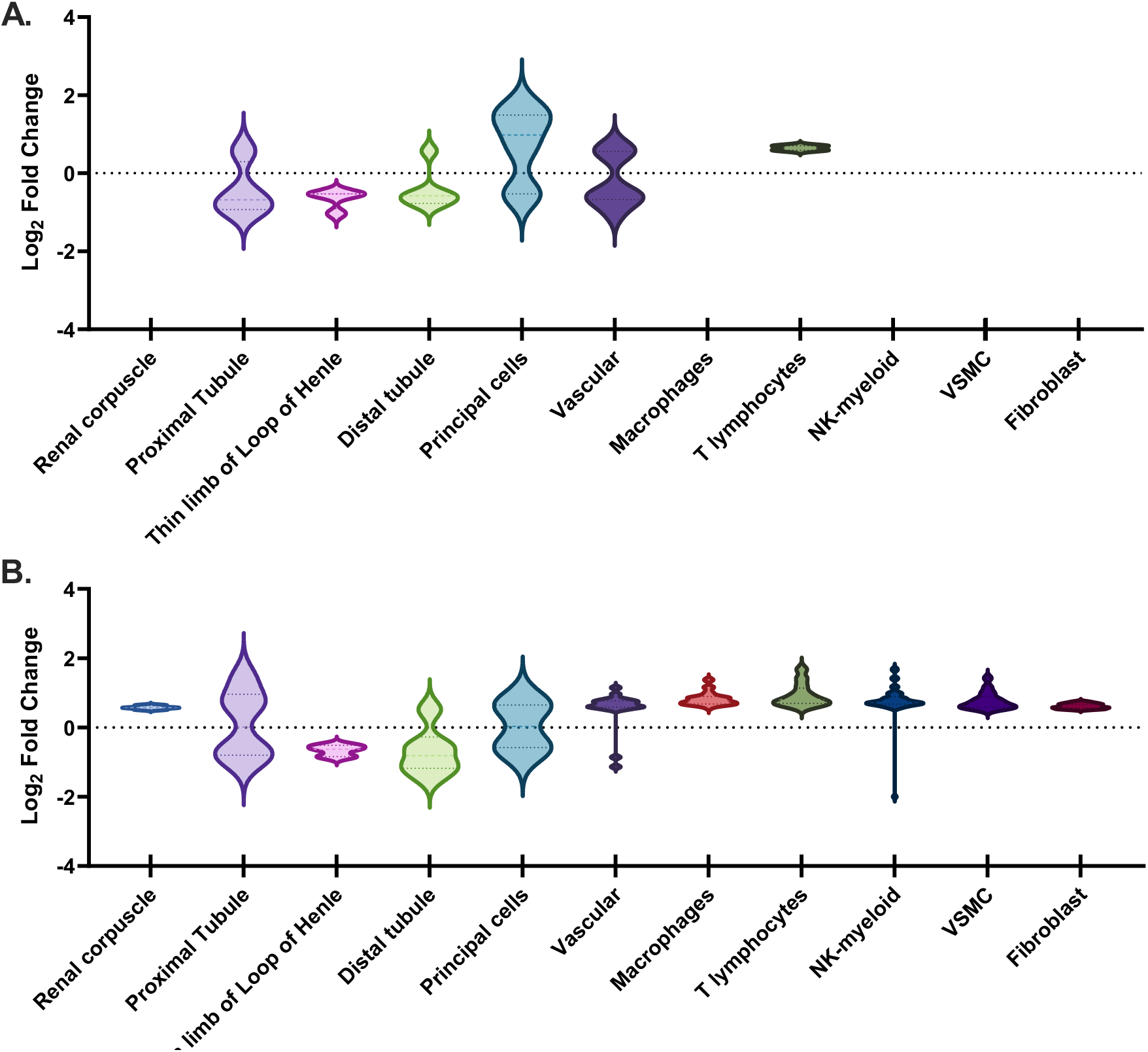
DEGs in Conv versus GF mice across renal cell types. Using data from Ransick, et al (12), single-cell transcriptomics of the mouse kidney reveals potential sex differences in gene expression. All DEGs characteristic of a particular renal cell type have been sorted under that cell type. Data are displayed as violin plots which show the distribution in log 2 fold changes for DEGs assigned to each cluster. We separately analyzed our data from females, to which 33 DEGs were assigned **(A)** and males, to which 176 DEGs were assigned **(B)**. Clusters without violin plot indicated cells with 1 of fewer DEGs. **NK-myeloid:** natural killer cell differentiation by myeloid lineage; **VSMC:** vascular smooth muscle cells.

## DISCUSSION

Although recent studies have explored the gut-kidney axis in health and disease (2, 18, 19), the role of host microbiota in renal gene regulation has not previously been examined. Here, we report that renal gene expression is altered by host microbiota, and that these changes are dependent on sex. Moreover, the majority of the genes regulated by microbes in kidney are not similarly regulated in the liver or large intestine, suggesting that host microbiota modulate gene expression in a tissue-dependent manner.

In our study, we conventionalized male and female GF mice using the same fecal slurry (made from a mix of male and female donors), and found that Conv males had a higher abundance of *Verrucomicrobia* than did Conv females. Previous data on sex differences in *Verrucomicrobia* are mixed: one study reported that female mice have a lower abundance of *Verrucomicrobia* compared to males (20), but another study found that males had a lower relative abundance of *Verrucomicrobia* compared to females (21). It may well be that *Verrucomicrobia* abundance depends on factors (food, environment, etc) which vary between different animal facilities.

Although we cannot rule out a role for sex differences in *Verrucomicrobia* to contribute to changes in host gene expression, this is unlikely to be solely responsible. Of note, the statistically significant difference in *Verrucomicrobia* was also observed at the species level, for *Akkermansia muciniphila (A. muciniphila*; p value = 0.005). A recent report observed that pre-treatment with *A*.*muciniphila* protected against lipopolysaccharide-induced acute kidney in mice (22), implying that this strain is physiologically important.

We also report here that the influence of the microbiota is tissue-specific. Previous studies identified a relationship between microbiome and host gene expression in the liver and large intestine (23, 24). However, we find that most genes regulated in the kidney are *not* regulated in the additional tissues sequenced. Utilizing GO Panther pathway analysis, we found that the top 100 female DEGs (Conv vs GF) unique to the kidney mapped to anion transport, regulation of neurogenesis, and rhythmic processes. For males, the top renal-specific pathways were mononuclear cell differentiation, negative regulation of immune system process, and positive regulation of response to external stimuli.

Our investigation revealed that only a small fraction of genes in kidney, liver, and large intestine were similarly regulated in both sexes. In agreement with this, previous studies have shown that the influence of microbiota on the host is often sex-specific (8, 25, 26). For example, Weger et al. reported that gut microbiota are required for sex-specific diurnal rhythms in gene expression (8). Similarly, a study utilizing the nonobese diabetic (NOD) mouse model (in which females develop type 1 diabetes at higher incidences than males) found that the sex difference in diabetes incidence is absent in GF animals(27). By comparing results from our bulk RNA-Seq to single cell RNA-Seq data from Ransick, et al (17), we mapped DEGs (Conv vs GF) to specific renal cell types. In addition to the sex bias that Ransick, et al previously reported in the proximal tubule, we observed a sex-bias in all other clusters.

Finally, our study revealed a small subset of genes which were commonly regulated across the three tissues we examined: *Per1* (in males), *Per2* (in females), *Mt1*, and *Mt2 (*in both males and females). Both *Per1* and *Per2* are associated with circadian rhythm; of note, additional genes related to circadian rhythm were found to be altered in Conv and GF mice, including *Cry1, Cry2, Ciart, and Bmal1*. In the kidney, *Cry1 and Cry2* were significantly upregulated in both male and female GF mice when compared to Conv. Furthermore, female GF mice exhibited increased renal expression (compared to male GF animals) in *Cry1, Cry2*, and *Ciart*. In the liver, *Ciart* was upregulated in GF males versus Conv males (log2fc = +4.03), and was not significantly changed in GF females versus Conv females (log2fc= -0.69, pvalue= 0.45). Finally, *Bmal1* was significantly upregulated in the livers of males (Conv vs GF), but not in the kidney. Previous studies have also highlighted the fact that germ-free mice have alterations in circadian biology genes (8). It may be that cyclical production of microbial metabolites such as short-chain fatty acids (which inhibit HDAC activity (28)) may modulate expression of circadian genes.

Furthermore, we report that *Mt1* and *Mt2* were significantly increased in the absence of gut microbiota in both sexes and in all three tissues. This is consistent with a report that *Mt2* was upregulated in the colon of GF mice using microarray (sex was not specified (29)), as well both *Mt1* and *Mt2* increased in the duodenum and the colon of GF mice (30). A notable difference between the Breton el al study and our study is that the former only examined females. We find that these changes are more pronounced in males (log2fold changes for Conv/GF in *Mt1* = +1.1, *Mt2* = +1.4, respectively) compared to females (*Mt1= +*0.6 and *Mt2*= +0.8).

In sum, we find that host microbiota modulate renal gene expression, and that this occurs in a sex-dependent and tissue-specific manner. Although host-microbiome interactions continue to be an evolving theme in physiology, microbiome-gene interactions remain largely unexplored. The human genome contains approximately 20,000 protein coding genes, whereas the microbial genome is approximately 100 times the number of genes in the human body (26). To this end, the gut microbiota has become increasingly recognized as the “second human genome” because its coding potential supersedes the coding potential of the human genome (26). Thus, in the future studies it will be important to expand upon our findings to understand the interaction between the gut microbiome and host transcriptomics, which can provide powerful insight into the physiological processes required to maintain host health.

## AUTHOR CONTRIBUTIONS

B.M. and J.L.P. developed the concept and designed the study; B.M. conducted the experiments and prepared the figures; B.M. and J.L.P. analyzed the data and interpreted the results; B.M. and J.L.P. drafted, edited, and approved the final version of the manuscript.

## ACKNOWLEDGEMENTS

We would like to thank the current and past members of the J.L. Pluznick laboratory for their insightful discussions. We would also like to thank Dr. Hua Ding, PhD and the animal care staff of the Johns Hopkins University Germ-Free facility; the JHU Single Cell and Transcriptomics Core for library prep and sequencing; Drs. Liliana Florea, PhD and Corina Antonescu, PhD for their help with RNA-Seq analysis; and finally, Dr. James Robert White, PhD of Resphera Biosciences for microbial sequencing analysis.

