## Supplementary Tables for "Commensal Microbiota Regulate Renal Gene Expression"

| FEMALE | | | | |
|---|---|---|---|---|
| Group | Cluster | Total DEGs | ↑ DEGs (increased in Conv vs. GF) | ↓ DEGs (decreased in Conv vs. GF) |
| Nephron | Renal corpuscle | 0 | -- | -- |
| | Proximal tubule | 4 | *Lpl* | *Cyp2e1,Inmt, Ttr* |
| | Thin limb of Loop of Henle | 4 | -- | *Cfap52,Hsp90aa1, Ifit3,Ifit3b* |
| | Distal tubule | 4 | *Lpl* | *Mt1,Mt2, Ptger3* |
| Ureteric epithelium | Principal cells | 7 | *Aldh1a3, Cbr2,Krt15,Krt5, Upk1b* | *Atp4a,Hsph1* |
| | Intercalated cells | 0 | -- | -- |
| Vascular | Vasculature | 8 | *Esm1,Lpl,Npr3* | *Fam167b,Ifit1,Isg15,Nostrin,Rgcc* |
| Immune cells | Macrophages | 1 | -- | *Snca* |
| | T lymphocytes | 2 | *Cd3e,Tcf7* | |
| | B lymphocytes | 0 | -- | -- |
| | NK-myeloid | 1 | -- | *Cd300a* |
| Interstitial cells | VSMC | 1 | -- | *Rarres2* |
| | Fibroblast | 1 | -- | *Rarres2* |

**Table S1. DEGs in Conv versus GF females within renal clusters.** Using data from Ransick, et al (12), table lists differentially expressed genes (DEGs) assigned to each renal cell type in female mice.

| MALE | | | | |
|---|---|---|---|---|
| Group | Cluster | Total DEGs | ↑ DEGs (increased in Conv vs. GF) | ↓ DEGs (decreased in Conv vs. GF) |
| Nephron | Renal corpuscle | 4 | *Fxyd1,Fhl2,Tmsb10,Rbp1* | -- |
| | Proximal tubule | 6 | *Cndp2,Prlr,Scd1* | *G6pc,Gsta2,Pck1* |
| | Thin limb of Loop of Henle | 3 | -- | *Arrdc3,Fst,Gdf15* |
| | Distal tubule | 4 | *Abca13* | *Mt1,Mt2, Ier3* |
| Ureteric epithelium | Principal cells | 2 | *Aldh1a3* | *Atp4a* |
| | Intercalated cells | 0 | -- | -- |
| Vascular | Vasculature | 20 | *Bst2,Ccrl1,Cd300lg,Ecscr,Esm1,Fbln5,Gng11,Ifi23,Ifi44,Ifit1,Irf7,Isg15,Lmo2,Marcks,Mmp11,Npr3,Rsad2,Sncg* | *Sgk1,Tmem252* |
| Immune cells | Macrophages | 36 | *Adgre1,Aif1,Bcl2a1b,C1qa,C1qb,C1qc,C3ar1,Cd52,Cd68,Cd72,Cd74,Ctss,Cx3cr1,Cybb,Fcer1g,Fcgr4,Fyb,H2-Aa,H2-Ab1,H2-Dma,H2Eb1,Laptm5,Lilra5,Lst1,Ly86,Mgl2,Mmp12,Ms4a7,Pid1,Pld4,Rgs10,Scimp,Slamf9,Spi1,Tyrobp,Wfdc17* | -- |
| | T lymphocytes | 24 | *Aw112010,Ccl5,Cd2,Cd3d,Cd3e,Cd3g,Cd52,Coro1a,Ctsw,Epsti1,Gimap3,Itgb2,Itgb7,Lat,Lck,Lsp1,Ltb,Nkg7,Ptprc,Rac2,Selplg,Tcf7,Thy1,Tmsb10* | |
| | B lymphocytes | 0 | -- | -- |
| | NK-myeloid | 46 | *Alox5ap,Aw112010,Bin2,Ccl5,Ccr2,Cd2,Cd209a,Cd300a,Cd52,Cd74,Clec10a,Clec12a,Coro1a,Ctss,Ctsw,Cybb,Fcer1g,Gimap3,H2-Aa,H2-Ab1,H2-Dma,H2-Dmb2,H2-Eb1,Itgal,Itgb2,Itgb7,Laptm5,Lck,Lsp1,Lst1,Mgl2,Ms4a6c,Nkg7,Pid1,Plbd1,Pld4,Ptprc,Rac2,Rnas26,Selplg,Slamf7,Spi1,Tmsb10,Tyrobp,Wfdc17* | *Nr4a1* |
| Interstitial cells | VSMC | 17 | *Cd2,Cd37,Cd52,Cd53,Cd79b,Coro1a,Des,Fxyd1,H2-Dmb2,H2-Ob,Lsp1,Ltb,Mgp,Pdgfrb,Rac2,Sncg,Tmsb10* | *Rarres2* |
| | Fibroblast | 14 | *Adamts5,Col1a2,Cxcl14,Des,Ecm1,Fbln5,Fhl2,Itga8,Lhfp,Mgp, Olfml3,Pdgfrb,Rcn3,Tcf21* | -- |

**Table S2. DEGs in Conv versus GF males within renal clusters.** Using data from Ransick, et al (12), table lists differentially expressed genes (DEGs) assigned to each renal cell type in male mice.
